# Genomic analysis revealed hotspots of genetic adaptation and risk of disappearance in the Brazilian goat populations

**DOI:** 10.1101/2024.06.03.597192

**Authors:** Francisco de A. Diniz Sobrinho, Ronaldo Cunha Coelho, Jeane de Oliveira Moura, Igor Ferreira do Nascimento, Miklos Maximiliano Bajay, Leonardo Castelo Branco Carvalho, Fábio Barros Britto, José Lindenberg Rocha Sarmento, Danielle Maria M. Ribeiro Azevedo, Salvatore Mastrangelo, Adriana Mello de Araújo

## Abstract

We accessed a 50K Illumina SNP genotype dataset from two important goat breeds of the Brazilian semi-arid region to analyze the abundance and length of runs of homozygosity (ROH). This analysis aims to elucidate the importance of adaptation history in the genome of the Brazilian goat populations and to measure genomic inbreeding. Heterozygosity-rich regions (HRR) or genome regions of high genetic variability provide clues about how diversity might be associated with increased fitness, avoiding deleterious homozygous alleles. Overall, 22,872 ROH were identified. The average number of ROH per individual ranged from 74.73 (Anglo-Nubian commercial breed) to 173.85 (Marota landrace). Analysis of the distribution of runs of homozygosity according to their size showed that, for both breeds, the majority of ROH were in the short (<2.0 Mb) category (65.6%). ROH-based inbreeding (F_ROH_) revealed low levels in Anglo-Nubian (0.0627) and high levels in Marota (0.1419), likely due to a reduction in effective population size over generations in the Marota landrace. We defined islands of ROH and HRR and identified common regions in the Marota goat, where genes related to various traits such as embryonic development, body growth, lipid homeostasis, and brain functions are located. These results indicate that such regions are associated with many traits and have therefore been under selective pressure in these goat breeds reared for different purposes.

## Introduction

Among livestock species, goats (*Capra hircus*) play a crucial role in economic development, significantly contributing to large-scale production, particularly of milk, and supporting rural subsistence by enhancing income for families in developing countries [1,2]. In the Brazilian context, farming activities involving small ruminants are divided between large producers and small breeders, with goats prominently adapted to the semi-arid tropics [3,4].

Goat landraces are generally more genetically diverse than commercial breeds, as they have been developed through a long history of breeding by processes markedly different from those used for commercial breeds. Several goat breeds adapted to the tropical climate of the semi-arid region of Brazil serve as valuable reservoirs of genes related to resistance to adverse conditions, parasites, and diseases, thus offering rich genetic diversity[5,6]. Therefore, identifying adapted genetic resources is essential to provide support and guidance in formulating policies to ensure the conservation and sustainable reproduction of this biological heritage. Hence, the sustainable management of farm animal genetic resources is an urgent and complex task to safeguard their significant contribution to the genetic heritage [7–11].

The introduction of large livestock in Brazilian history can be divided into the eras of colonialism and the 1950s green revolution: the Marota landrace (Portuguese naturalized) and the Anglo-Nubian (commercial) are two significant representatives of these periods. Marota goats inhabit the semi-arid region; they are a distinct local type with a white coat, small size, and hardy nature, maintained by government institutions and partners in northeastern Brazil. This population constitutes a valuable reservoir of genetic diversity, as they possess genes linked to minimizing negative effects associated with climate and environmental changes, as well as resistance to parasites and diseases, attributable to their evolutionary history of adaptability to harsh environmental conditions[3,5,12]. Despite its long history in Brazil, the genetic diversity and population structure of the Marota landrace remain poorly understood.

Autozygosity, the homozygous state of identical-by-descent (IBD) alleles, can result from several phenomena. An increase in inbreeding (F) leads to various negative effects, such as a reduction in genetic variance, decreased individual performance (inbreeding depression), and lower population viability [13,14]. Currently, among various methods, F estimated from ROH is considered the most effective[15,16]. ROH can also be used to identify genomic regions potentially under selection and involved in defining population-specific traits[14,16,17].

Considering gene recombination, genomic regions with a large amount of homozygosity are presumably the result of selection, leading to favorable alleles in surrounding regions. Thus, ROH analysis has been employed to explore signatures of selection in various species, including both ruminants and non- ruminants [18–21].

The pattern of heterozygosity-rich regions (HRR) in the genome can also offer insights into population structure and demographic history. HRR islands are previously associated with heterozygous advantage. In statistical analysis of consecutive runs, HRR are linked with adaptive traits such as response to heat stress, immune response, survival rate, fertility, and other fitness-related traits [16,22].

Considering the evolutionary factors that drive population genomic structure, the present study provides a comprehensive genome-wide population analysis, characterizing patterns of ROH and HRR in two goat breeds. As such, this work significantly contributes to understanding genetic diversity and evolutionary processes in Brazilian goat populations, highlighting the importance of F_ROH_ for the conservation and sustainable management of these genetic resources.

## Methods

### Ethics statement

The database for this study, titled “Population genomics, genetic diversity, and gene introgression in naturally adapted goat conservation flocks in Brazil,” was approved by the Ethics and Research Committee of the Federal University of Piauí, adhering to the established guidelines, with registration number 058/14.

### Location of sampled data

The state of Piauí, located in the northeast region of Brazil, borders the states of Maranhão, Ceará, and Pernambuco (Fig 1). It spans an area of 251,529 km^2^ and harbors the third-largest goat population in Brazil (IBGE, 2017).

**Fig 1.**
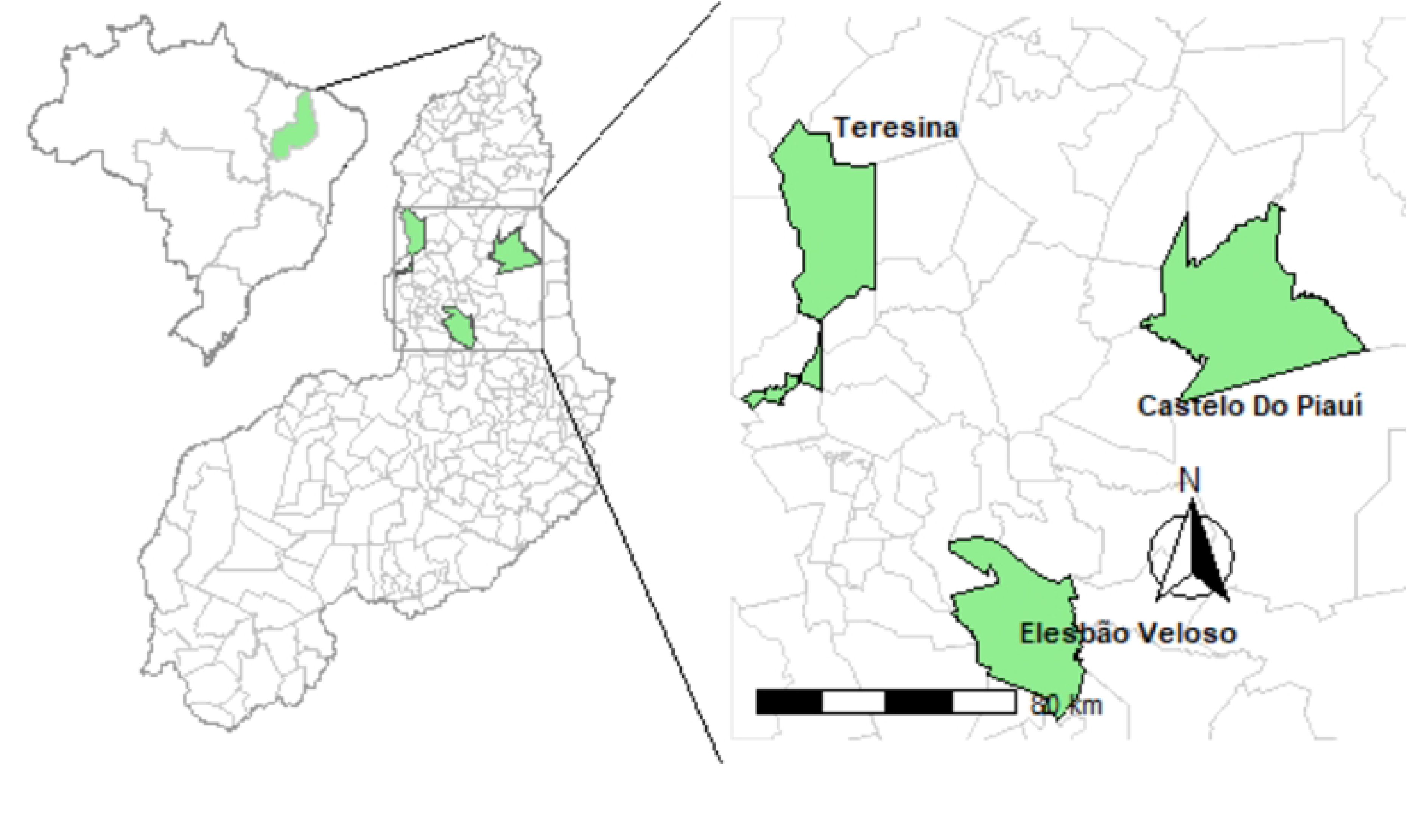
Geographical and phenotypic distribution of two goat populations in Piauí, Brazil.

The data used in this study included 192 animals from two distinct breeds found in the semi-arid biome: the rare landrace Marota (n = 86), and the standardized breed Anglo-Nubian (n = 106). Individuals from the Anglo-Nubian breed were acquired from the municipality of Teresina, PI, Brazil (05°02’39.95” S, 42°47’03.70” W), while those from the Marota landrace were provided the National Goat Conservation Program at Embrapa (05°19’20” S, 41°33’09” W) and a private farm in Elesbão Veloso, PI, Brazil (06°12’07” S, 42°08’25” W).

### Genotyping

Genotyping was performed using the 50K Illumina Goat Bead Chip, which contains 53,347 evenly spaced SNPs. This chip employs Illumina™ Infinium technology and the iScan platform. The genotyping protocol adhered to the guidelines provided by the manufacturer, available at www.illumina.com[3,4].

### Quality control

The quality of the genotypic data was assessed using PLINK version 1.9 [23]. The chromosomal coordinates of SNPs were aligned to the ARS1 genome assembly (https://www.ncbi.nlm.nih.gov/datasets/genome/GCA_000317765.1/). Markers assigned to unmapped contigs and to sex chromosomes were excluded from further analysis. Quality control metrics included a call-rate threshold of ≥0.95 [16,24], yielding a final dataset of 43,133 SNPs.

### Detection of ROH and HRR

The detectRUNS package, as proposed by Biscarini et al. [27] for the R environment, was used to detect both ROH and HRR using the sliding window and consecutive runs methods [23,25].

The following sliding window constraints were imposed for ROH detection: (i) window size of 20 SNPs across the genome (ii) the minimum number of SNPs included in a sliding window was set at 20 (minSNP); (iii) the maximum number of heterozygotes allowed in a run was 1 (maxOppWindow); (iii) the number of missing SNPs allowed per window was 1 (maxMissWindow). Run-related parameters included: (i) the maximum gap between consecutive SNPs was set to 250 kb (maxGap = 250,000 bps); (ii) the minimum ROH length was established at 1000 kb (minLengthBps = 1,000,000); (iii) the minimum density of one SNP every 70 kb (minDensity-1/70); (iv) the proportion of homozygous overlapping windows was set at 0.05[16,25–31].

The percentage of chromosomes covered by ROH was estimated following the method proposed by Al-Mamun et al. [33]. We calculated the frequency and average length of the ROH for each length category (1–2, 2–4, 4–8, 8–16, and >16 Mb). Additionally, we determined the total number of ROH identified in each length category for each individual of each breed.

The genomic inbreeding coefficient (F_ROH_) for each breed was calculated using the following method (Equation 1), as described by McQuillan et al. [32]:

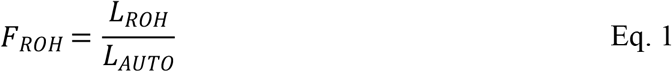

where **L**_**ROH**_ represents the total length of ROH of each individual in the **genome**, and **L**_**AUTO**_ is the length of the autosomal genome of the goat, set at 2,464.80 Mb[32–34].

To identify the genomic regions that were most commonly associated with ROH, the SNP frequencies (%) within detected ROH were assessed for each breed and plotted against SNP positions across autosomes. ROH islands in a breed were defined as those regions with a frequency ≥ 50%[35–39].

HRR detection employed the consecutive runs (CS) method, with the following parameters: (i) a minimum of 10 SNPs in a run; (ii) a maximum of five homozygous SNPs in a run; (iii) a maximum of five missing SNPs in a run; (iv) the minimum HRR length was set to 1 Mb; (v) the maximum gap between consecutive SNPs was set to 1 Mb [25]. HRR segments are predicted to be less frequent and shorter (<1 Mb) than ROH segments. The primary criterion for their identification is the number of allowed homozygous SNPs within a segment[18,25,40]. HRR must present a frequency of 45% in individuals in the population[14].

## Results

### Detection of ROH

Overall, 22,872 ROH were identified across 192 individuals. Figure 2 shows the percentage of chromosomes covered by ROH and the number of ROH per chromosome. The highest coverage by ROH was observed on chromosome 1. The number of ROH per chromosome displayed a same pattern with the highest numbers found for the first chromosome, a number that tended to decrease with chromosome length.

**Fig 2.**
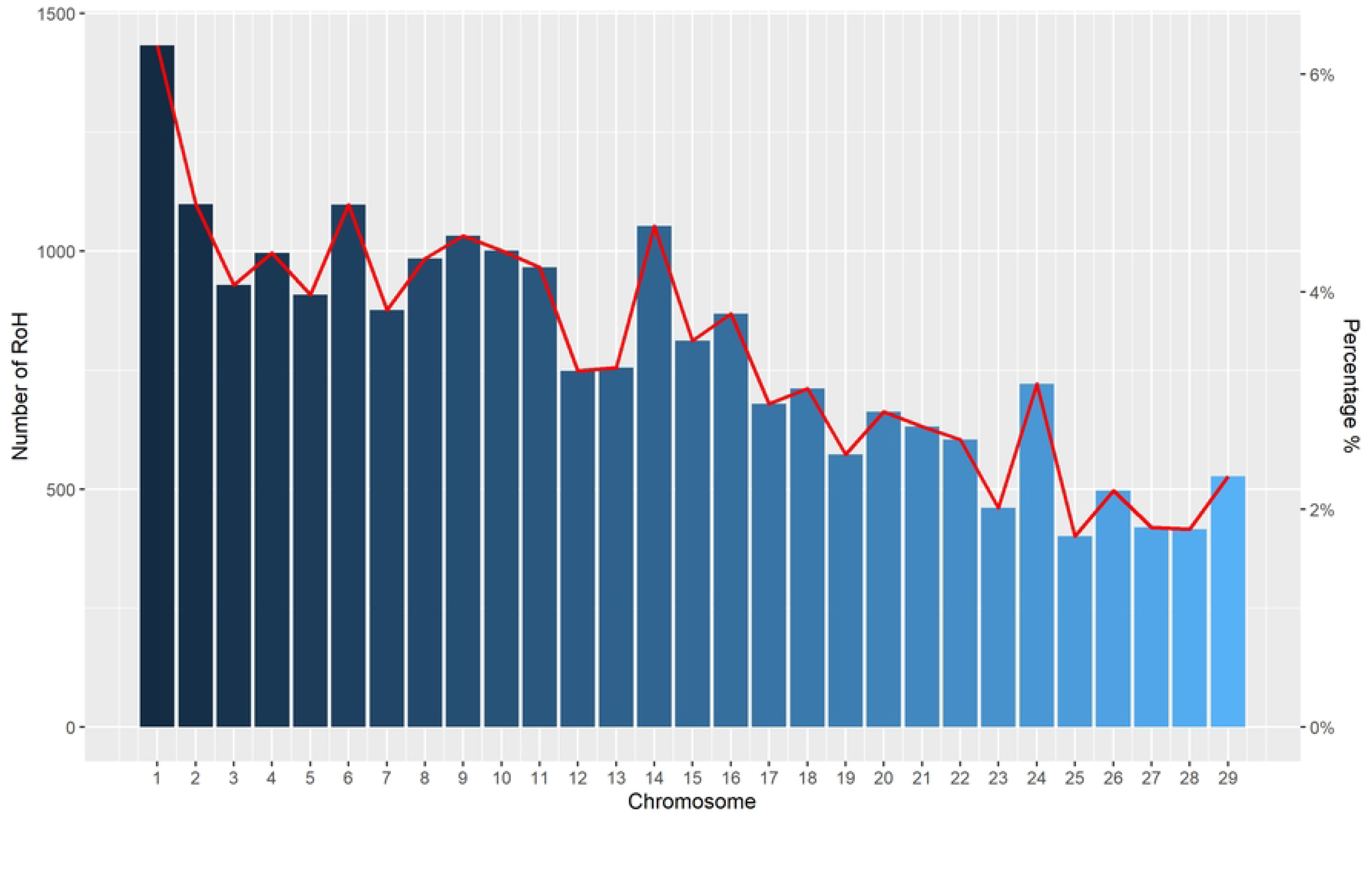
Number of ROH per chromosome (bars) and average percentage of each chromosome covered by ROH (lines) for all goats.

The ROH segments varied in length from 1.0 Mb to 14.55 Mb in the traditional Marota breed and from 1.0 Mb to 12.87 Mb in the Anglo-Nubian breed. The number of ROH ranged from 4 to 270 in Anglo-Nubian goats and from 56 to 269 in Marota goats (Table 1).

**Table 1.** ROH length and number in goat populations from the Brazilian semi-arid region.

| Breed | n | ROH length |  |  | ROH number |  |  |
| --- | --- | --- | --- | --- | --- | --- | --- |
|  |  | Mean | Min | Max | Mean | Min | Max |
| Marota | 86 | 2.01 ± 1.13 | 1.00 | 14.55 | 173.85 ± 49.46 | 56 | 269 |
| Anglo-Nubian | 106 | 2.07 ± 1.12 | 1.00 | 12.87 | 74.73 ± 45.9 | 4 | 270 |

*n* = number of individuals; *Min* = minimum values observed in estimations of individual animals within each breed; *Max* = maximum values observed within each breed.

### Genomic inbreeding (F_ROH_) coefficients

The average genomic inbreeding coefficient (F_ROH_) in the Anglo-Nubian breed was 0.0627, with a variation of 0.003 ± 0.323. Higher values were found in the Marota landrace, reaching 0.1419, with a variation of 0.040 ± 0.283 (Fig 3).

**Fig 3.**
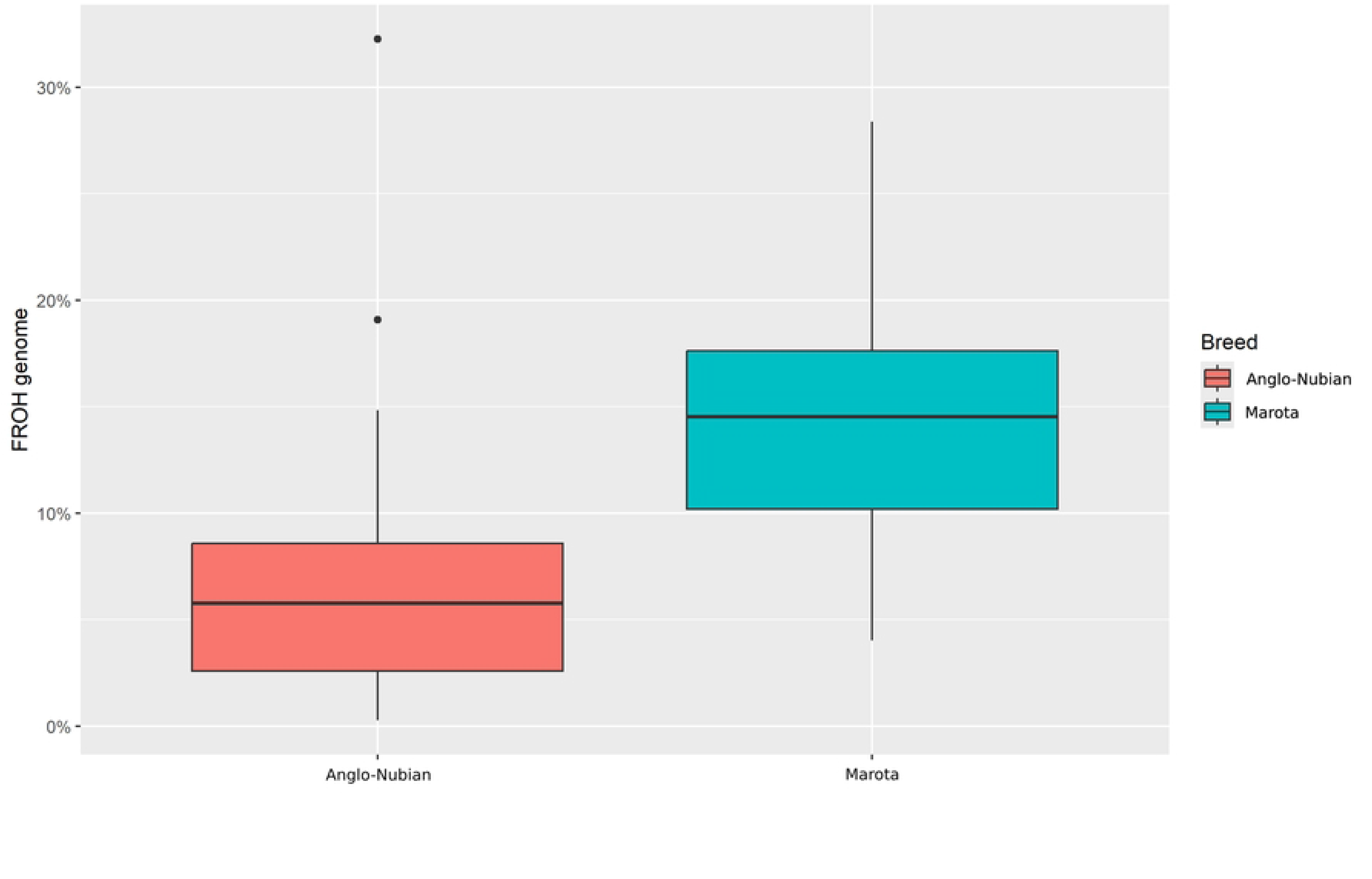
Boxplot of F_ROH_ distribution within the Anglo-Nubian and Marota goat breeds.

To investigate recent and past inbreeding in each breed, the distribution of different ROH size classes was established (Table 2). All breeds displayed more ROH in the 1-2 Mb class than in other classes. A higher total number of ROH occurred in the Marota landrace (14,951), but the Anglo-Nubian breed had approximately the same percentage distribution of ROH.

**Table 2.** Summary of number of runs of homozygosity (ROH) by size class in Marota and Anglo-Nubian goats.

| Class (Mb) | Marota |  | Anglo-Nubian |  |
| --- | --- | --- | --- | --- |
| 1 - 2 | 9,813 | 65.60% | 4,260 | 62.60% |
| 2 - 4 | 4,962 | 28.50% | 2,473 | 31.20% |
| 4 - 8 | 818 | 5.50% | 456 | 5.80% |
| 8 - 16 | 60 | 0.40% | 30 | 0.40% |
| > 16 | 0 | 0.00% | 0 | 0.00% |
| Total | 14,951 | 100% | 7,921 | 100% |

### Run of homozygosity islands

To identify genomic hotspot regions for selection and/or conservation, the frequency of SNPs within ROH across the autosomes was plotted (Fig 4). Generally, adjacent SNPs with an ROH frequency ≥ 50% and a minimum of **14** SNPs above the adopted threshold form genomic regions called ROH islands.

**Fig 4.**
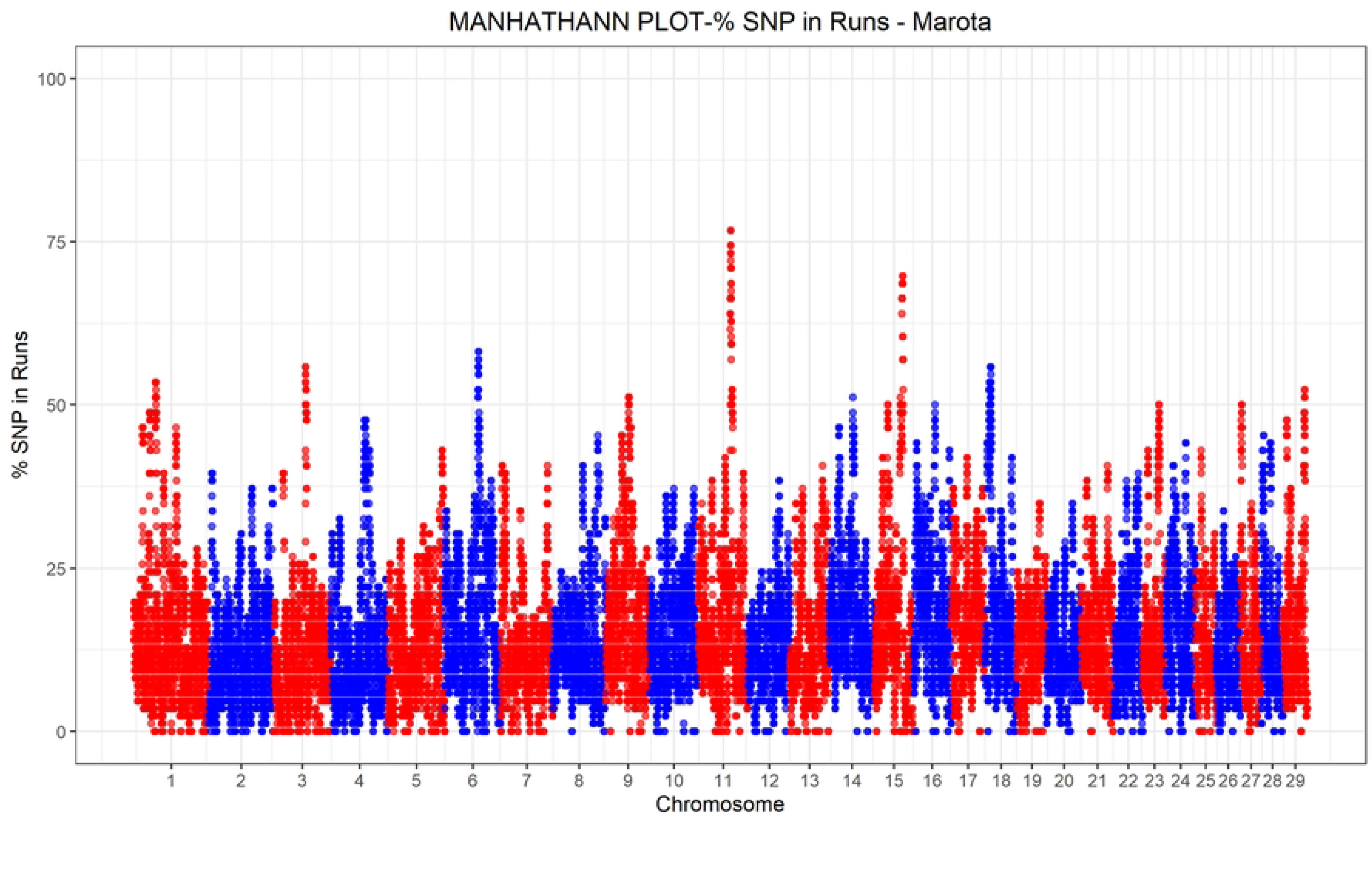
Manhattan plot of the proportion of times each SNP falls within a ROH in the Marota goat landrace.

Applying a threshold of 50% on the Manhattan plot, seven regions exhibiting a high occurrence of ROH were identified in the Marota landrace (Figure 4). No ROH islands met the 50% threshold in the Anglo-Nubian Manhattan plot. A map of these target genomic regions is provided in Table 3. Autosomes 01, 06, 11, 15, and 18 consistently showed high ROH occurrence.

**Table 3.**
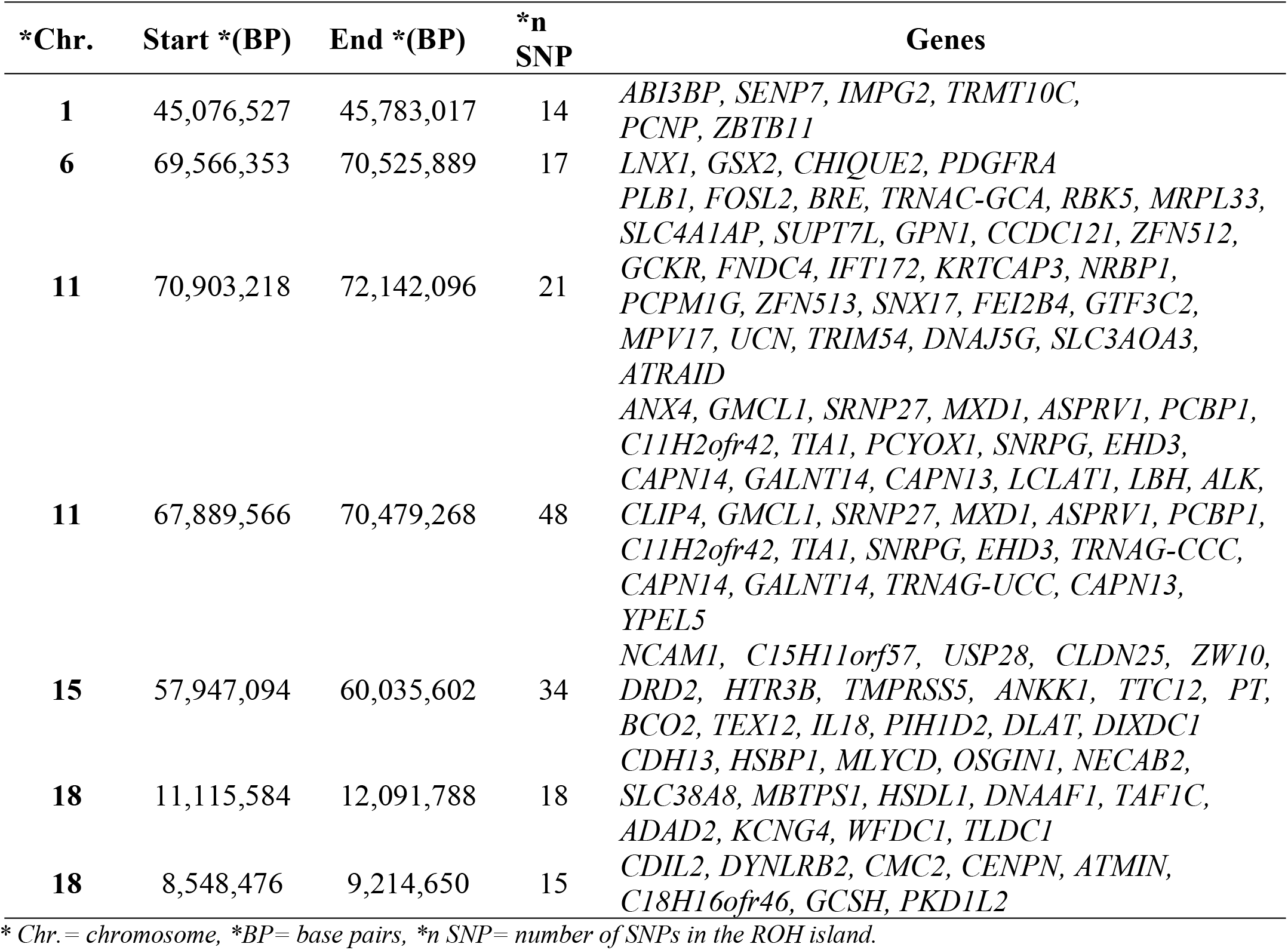
Gene annotation of runs of homozygosity (ROH islands) found in the autosomal genome of the Marota goat population.

### Heterozygosity-rich regions

The HRR analysis, utilizing consecutive runs (CR), identified 17,278 spots in total. Table 4 provides a descriptive summary of the length and abundance of HRR. Based on the analysis, regions identified in less than 45% of animals were deemed HRR islands and were subsequently scrutinized for candidate genes and pathways. Specific HRR islands discovered in the Marota and Anglo-Nubian goat breeds are listed in Table 5.

**Table 4.** HRR length and number in goat populations from the Brazilian semi-arid region.

| Breed | n | HRR length |  |  | HRR number |  |  |
| --- | --- | --- | --- | --- | --- | --- | --- |
|  |  | Mean | Min | Max | Mean | Min | Max |
| Marota | 86 | 1.18 ± 0.2 | 1.00 | 2.98 | 67.24 ± 15.49 | 41 | 360 |
| Anglo-Nubian | 106 | 1.18 ± 0.18 | 1.00 | 2.63 | 108.44 ± 34.59 | 48 | 108 |

*n* = number of individuals; *Min* = minimum values observed in estimations of individual animals within each breed; *Max* = maximum values observed within each breed.

**Table 5.**
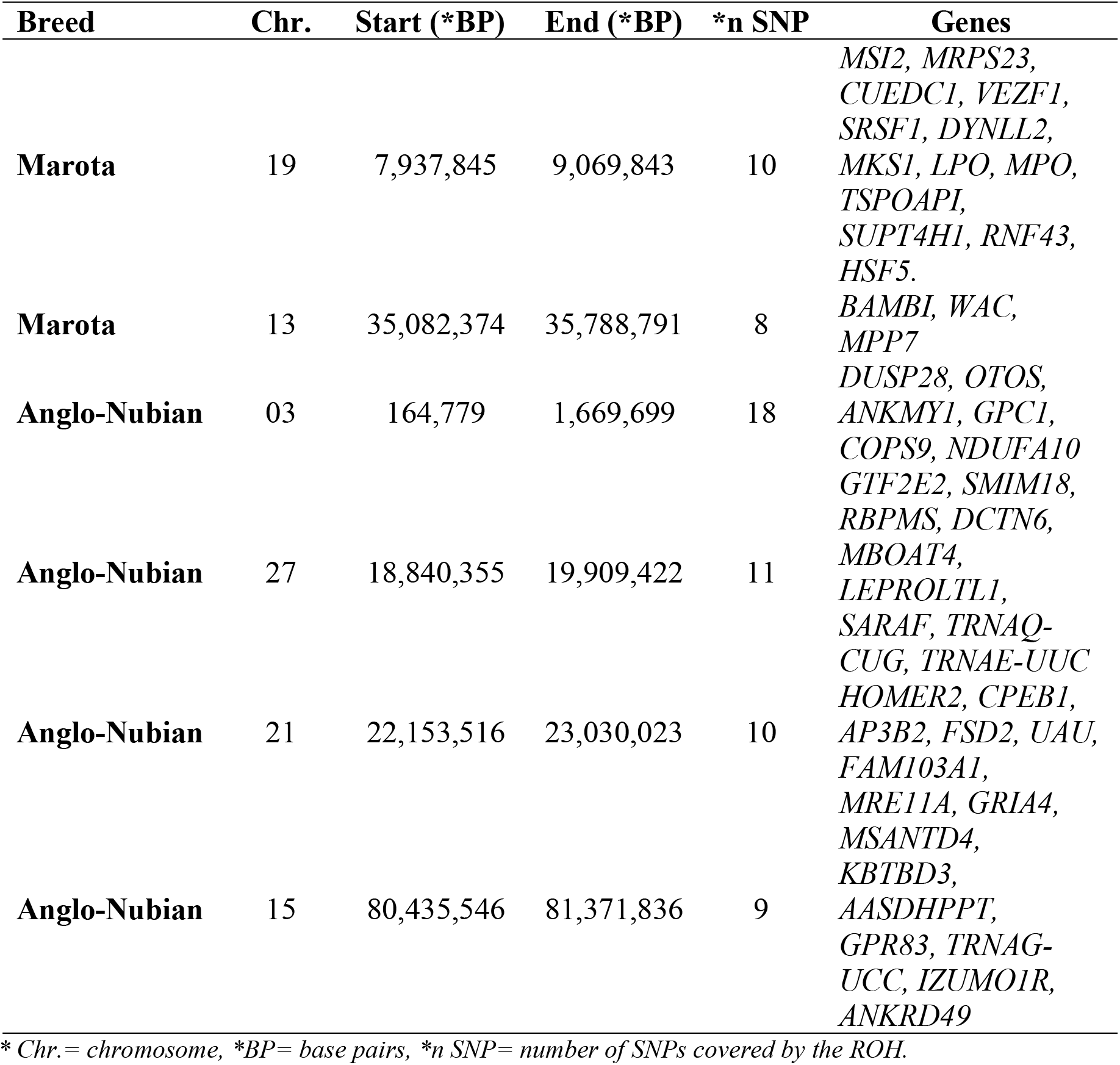
Gene annotation of the heterozygosity-rich regions (HRR islands) found in the autosomal genome of Marota and Anglo-Nubian goats.

## Discussion

### Homozygosity patterns (ROH)

The analysis of homozygosity patterns (ROH) has emerged as an informative tool for elucidating the demographic history of populations. It provides precise information on genetic diversity, and the assessment of ROH lengths enables the detection of both past and recent genomic inbreeding in animals [41–43].

The number of ROH in these breeds is higher compared to Turkish (60), Russian (approximately 73), and Central Asian (90) indigenous breeds[38,44]. Compared to standard breeds in America, Anglo-Nubian goats exhibit lower ROH values than those reported in related studies (133)[45].

In general, variations in the number and distribution of ROH may reflect historical differences in genomic structures. Highly selected breeds often show a greater abundance of ROH and broader coverage compared to local breeds [15,45–47]. In our study, the Marota landrace is distinguished by its high number of ROH, contrasting with the Anglo-Nubian commercial breed. Bertolini et al. [45] noted an increased trend of homozygosity in local breeds, attributed to their small population sizes and geographical isolation.

### Genomic inbreeding of homozygosity (F_ROH_)

This study examines the level of genomic inbreeding in goat breeds in the Brazilian semi-arid region. Understanding the level of endogamy in a population has practical applications in management and enhances the understanding of relationships between individuals. It helps estimate inbreeding levels, correct errors often found in family trees, and identify undocumented ancestors in pedigrees. This method is particularly beneficial for studying genetic variation in small, at-risk populations[48,49].

The average F_ROH_ in Brazilian Marota goats (0.1419) is higher than that of other globally conserved goat breeds (F_ROH_ = 0.12). In contrast, Anglo-Nubian goats show lower F_ROH_ values (0.0627) and exhibit a lower degree of inbreeding compared to several other breeds[50]. The higher genome inbreeding in the Marota goats may result in closer related individuals, leading to a significant prevalence of recessive genes in homozygotes, increased inbreeding, and genetic disorders. Therefore, closely monitoring the population is essential to prevent the loss of genetic resources and minimize the negative effects of harmful mutations, inbreeding depression, and loss of genetic diversity[42,45,51,52].

A significant prevalence of short ROH segments was observed. Specifically, 21,508 segments were identified in the smallest size category (<4 Mb), representing approximately 94% of the total ROH values obtained in the genomes of the Marota and Anglo-Nubian breeds. Comparing ROH results is a challenging task due to the variety of criteria used in different studies. Our research revealed that in the Marota and Anglo-Nubian breeds, the majority of ROH are of short size. Hence, our findings suggest that these animals experienced long-term inbreeding [18,51–53].

Deniskova et al. [41] reported similar results for *ROH*<4*MB* using a 50K panel for Russian goat. The emphasis on the prevalence of short-length ROH, particularly in Marota goats, underscores the need for management strategies and conservation efforts to mitigate inbreeding and safeguard adaptive traits, thereby ensuring genetic diversity in local goat populations[42,54].

### Runs of homozygosity and heterozygosity-rich regions

Livestock farming is a human practice influenced by various environmental factors. When environmental conditions are unfavorable, animals can adapt in diverse ways to cope with stress. Thus, biological adaptation is essential to ensure that animals remain healthy and productive. Within their DNA, it is possible to find markers of natural selection that emerge in response to environmental pressures, identifiable through genomic and bioinformatics techniques[18,54,55].

In this context, goat breeds from the Brazilian semi-arid region demonstrate adaptive evolution aimed at survival, reproduction, and production across different ecological zones. It is likely that their genomes have developed unique genetic characteristics, evolving to adapt to remarkably heterogeneous environments[3,6]. For this reason, it is expected that both artificial and natural selection would influence animal genomic diversity, leaving behind either positive or deleterious adaptive signatures[52,56].

Analysis of ROH and HRR in the Marota and Anglo-Nubian breeds has revealed markers potentially under selection for adaptation to the semi-arid environment of Brazil. Our discoveries have led to the identification of target genes related to adaptation, specifically concerning the immune/inflammatory response, energy homeostasis, reproductive and production traits, and heat stress[18,22,40,54,55,57].

Research into the specific functions of genes has unveiled an intricate panorama of biological processes, wherein genes play distinct and essential roles in cellular functions, contributing to a wide range of biological processes. For example, the *GMCL1* and *TEX12* genes are known to be involved in the regulation and/or participation of reproductive processes, specifically in sperm production[9,18,55,56]. The *Zbtb11* gene, with its integrase-like HHCC zinc finger, serves as a crucial regulator in neutrophil differentiation, contributing significantly to the immune defense system. Similarly, the *IL18* gene expresses the IL-18 protein, a pro-inflammatory cytokine pivotal for immune response regulation, activating immune system cells like T lymphocytes and natural killer (NK) cells, thereby enhancing the production of inflammatory cytokines[58,59].

The *LBH* and *YPEL5* genes play specific roles in embryonic development, RNA processing, and cell cycle regulation, thereby influencing cell growth [60]. The data indicate that certain genes, subject to natural selection, are involved in a wide range of cellular metabolic processes, including protein cleavage and lipid homeostasis. These genes include *PCYOX1, CAPN14, GALNT14, TRNAG*-*UCC, CAPN13*, and *LCLAT1* [61–64]. Genes such as *CLDN25* and *ZW10* actively participate in regulating cellular permeability and chromosomal segregation during cell division and may play roles in cell growth[65,66].

Heterozygosity-rich regions present a new concept introduced by Williams et al. [44], being much less characterized than ROH in livestock. They are identified as heterozygous genomic regions potentially associated with disease resistance, immunity, and adaptive processes, serving as a valuable genetic reservoir and providing additional insights into goat genomes [24].

Heterozygosity loci are clustered in islands throughout the genome and are significantly rarer and shorter compared to ROH [22]. Several studies have reported divergent means of HRR per chromosome [18,22,67]. Variations in methods, animal populations, and SNP arrays may account for differences from our results, a trend similarly observed by Chessari et al. [24].

After applying a threshold of 45%, there were 66 SNPs. These SNPs formed six regions across six chromosomes, detailed in Table 5. They are located on chromosomes 3, 13, 15, 19, 21, and 27, determined by a single region. In these regions, we identified 46 gene. Based on genetic functions obtained from http://www.genecards.org (last accessed on July 4, 2024), we conclude that the genes found in the HRR are important in adaptive biological processes.

Of the 46 gene annotations, three genes with known and well-described functions are particularly notable. The *MPO* gene, located on chromosome 19, is responsible for encoding the enzyme myeloperoxidase. In humans, myeloperoxidase is involved in generating free radicals and hypochlorite, released by neutrophils during the inflammatory response to bacterial infections[68].

Similarly, the *LPO* gene encodes the enzyme lactoperoxidase, which plays a fundamental role in immune defense against bacterial infections by catalyzing the production of HOSCN from H_2_O_2_ and SCN^-^, a component of the antibacterial defense system present in the respiratory tract [69].

The *WAC* gene is a vital component of the molecular mechanism that coordinates the cell’s response to genotoxic stress, ensuring genome integrity and maintaining cellular homeostasis through transcriptional regulation of p53 target genes[70].

Our findings are in line with previous research involving Russian cattle, particularly the identification of a single candidate region on chromosome 15 of the Anglo-Nubian goat. Several genes are located within this region, namely MSANTD4 and GRIA4. Both genes are potential contributors to the climate stress resistance phenotype, given their indirect functions in heat shock response (MSANTD4) and body thermoregulation (GRIA4)[71].

## Conclusion

Genomic diversity and ROH parameters reveal well-defined differences that correspond to the geographical distribution of goats, as well as their breeding history and population size.

The ROH models indicated low levels of crossbreeding and low gene flow between the Anglo- Nubian breed and the Brazilian landrace. However, high F_ROH_ values in the Marota goats may seriously affect their overall biological fitness. High inbreeding levels are responsible for the fixation of genomic regions carrying deleterious mutations, which can push the population toward an extinction vortex.

Genes found on ROH islands and HRR are associated with relevant pathways of environmental adaptation. Thus, the presence of ROH and HRR islands in the genome of landraces highlights the evolutionary forces that drive adaptation.

Finally, ROH and HRR approaches underscores the importance of utilizing applied genome marker- based data to mitigate the loss of diversity in the future and to inform the development of more effective breeding and conservation programs.

## Authors’ contribution

### Conceptualization

Francisco de Assis diniz Sobrinho, Miklos Maximiliano Bajay and Adriana Mello de Araújo.

### Methodology

Francisco de Assis Diniz Sobrinho, Salvatore Mastrangelo, Igor Ferreira do Nascimento, Miklos Maximiliano Bajay, and Leonardo Castelo Branco Carvalho

### Validation

Fábio Barros Britto, Adriana Mello de Araújo, and Salvatore Mastrangelo

### Formal analysis

Francisco de Assis Diniz Sobrinho, Ronaldo Cunha Coelho, Igor Ferreira do Nascimento, Miklos Maximiliano Bajay, and Leonardo Castelo Branco Carvalho.

### Data curation

Jeane Oliveira Moura, Danielle Azevedo and José, and Lindemberg Sarmento.

### Writing - preparation of original draft

Francisco de Assis Diniz Sobrinho

### Writing - revision and editing

Adriana Mello de Araújo, Salvatore Mastrangelo, Miklos Maximiliano Bajay, Leonardo Castelo Branco Carvalho, Fábio Barros Britto, and Danielle Azevedo

### Supervision

Miklos Maximiliano Bajay and Adriana Mello de Araújo.

## Financing

Not applicable

## Statement by the Institutional Review Board

Not applicable.

## Informed Consent Statement

Not applicable.

## Data Availability Statement

Data available on request.

## Acknowledgments

We thank the Foundation of Support to Research of Piauí (FAPEPI).

